# Resolving the Interactions between Class 3 Semaphorin Receptors in Live Cells

**DOI:** 10.1101/2021.02.18.431813

**Authors:** Shaun M. Christie, Jing Hao, Erin Tracy, Matthias Buck, Jennifer S. Yu, Adam W. Smith

## Abstract

The plexin/neuropilin/semaphorin family of proteins is involved with tissue patterning in the developing embryo. These proteins play roles in cell migration and adhesion, but are also important in disease, including cancer angiogenesis and metastasis. While some structures of the soluble domains of these proteins have been determined, the conformations of full-length receptor complexes are just beginning to be studied, especially within the context of the cell plasma membrane. Pulsed-interleaved excitation fluorescence cross-correlation spectroscopy (PIE-FCCS) allows direct insight to the formation of protein-protein interactions in the membrane of live cells. Here we investigated the homodimerization of neuropilin-1, Plexin A2, Plexin A4, and Plexin D1. Consistent with previous studies, we found that neuropilin-1, Plexin A2 and Plexin A4 are dimers in the absence of exogenous ligand. Plexin D1, on the other hand, was monomeric under similar conditions, which had not been previously reported. We also found that Plexin A2 and A4 assemble into a heteromeric complex. Stimulation with Semaphorin 3A or Semaphorin 3C ligand neither disrupts nor enhances the dimerization of the receptors when they are expressed alone, suggesting that activation involves a conformational change rather than a shift in the monomer-dimer equilibrium. However, upon stimulation with Semaphorin 3C, Plexin D1 and neuropilin-1 form a heteromeric complex, while Semaphorin 3A does not induce a stable complex with these receptors. This analysis of interactions by PIE-FCCS provides a complementary approach to the existing structural and biochemical data that will aid in the development of new therapeutic strategies to target these receptors during disease.

## Introduction

The semaphorins are a large family of secreted and transmembrane ligands that regulate cell morphology and motility during development in a broad range of tissues.^1^ Twenty members of this ligand family are found in vertebrates, where they are categorized by homology into classes 3-7.^2^ The plexin family are type I transmembrane receptors and act as the main binding partner for these ligands at the cell membrane. Nine plexins are found in vertebrates, grouped in class A-D based on homology.^2–3^ Secreted class 3 semaphorins require an additional receptor moiety, neuropilin-1 or neuropilin-2 (Nrp1 and Nrp2), which have no intrinsic enzymatic activity, but create a holoreceptor complex with plexin to promote signalling.^2, 4–6^ Complex formation is a necessary part of their signaling activity, yet a profile of these interactions is still lacking due to the difficulty of working with membrane proteins in their native environment. Studies of plexin/neuropilin/semaphorin have mostly focused on biochemical data or structures of soluble domains of the proteins to establish protein-protein interactions while receptor interaction in the live cell environment has not been fully explored.^7^ Using pulsed interleaved excitation fluorescence cross-correlation spectroscopy (PIE-FCCS) we are able to complement the cellular data and other biophysical methods in order to understand a broader range of interactions and their likely role in plexin mediated signaling.^8^

Of the class 3 semaphorins, Semaphorin 3A is the most well studied for its function as a chemorepellent in axon guidance and growth cone collapse, where deletion can lead to excessive axonal branching.^9–14^ However, this ligand has broad expression across tissues where it plays multiple functional roles, such as vessel branching of the developing cardiovascular system, lungs, and kidneys.^15–18^ Another class 3 semaphorin, Semaphorin 3C, is expressed in the developing nervous system where it acts as a repulsive cue to guide tissue borders.^19^ Signaling by this ligand is also required for cardiovascular and lung development, with knockout mice, in some genetic backgrounds, unlikely to survive past the first few days.^16, 20–23^ While these ligands are an integral part of development, changes to their expression can lead to disease states. Dysregulation of Semaphorin 3A or Semaphorin 3C this has been implied for cardiovascular disease and various cancers.^21, 24–27^

Signaling is initiated when semaphorins bind to plexin receptors. Semaphorin 3C regulates downstream signaling in the presence of Nrp1,^20^ Plexin B1,^28^ Plexin A2, or Plexin D1.^29^ Various co-IP experiments have suggested the formation of complexes between multiple combinations of these receptors.^30–32^ However, co-IP may not be completely accurate due to the removal of proteins from the cell membrane environment and loss of inhibitory conformations.^33^ The goal of this work was to determine which receptors interact prior to and following Semaphorin 3A and Semaphorin 3C stimulation in the membrane of live cells using PIE-FCCS.

Quantifying the interactions between membrane proteins is experimentally challenging, and only a few of the plexins have been investigated with quantitative biophysical methods. Our laboratory first reported the ligand-independent homodimerization of Plexin A4 using PIE-FCCS.^34^ In that study, we found that deletion of the sema domain abrogated homodimerization. Later, the structure of this interaction was resolved for Plexin A4, as well as Plexin A2 and Plexin A1, by Kong *et al.* using X-ray crystallography and verified with FLIM-FRET.^35^ Nrp1 and Plexin A2 dimers have also been investigated with co-IP and quantitative FRET assays. These studies reported that Nrp1 forms small multimers in its basal state, but transitions to dimers following ligand stimulation.^36–37^ As noted, Semaphorin 3A and Semaphorin 3C require Nrp1 in order to induce signaling through complex formation with plexin receptors, such as with Plexin A2.^32, 38^ A 7.0 Å low/medium resolution structure for the tripartite interaction of Semaphorin 3A, Plexin A2, and Nrp1 ECDs has been solved where the complex suggests a 2:2:2 stoichiometry with Nrp1 acting as the bridge between Semaphorin 3A and Plexin A2.^39^ A schematic for this type of signaling complex is shown in Figure 1. In addition, signal propagation through Plexin A4-Nrp1 complexes is supported by cell collapse and alkaline phosphatase (AP) binding assays.^10, 40–41^

**Figure 1.**
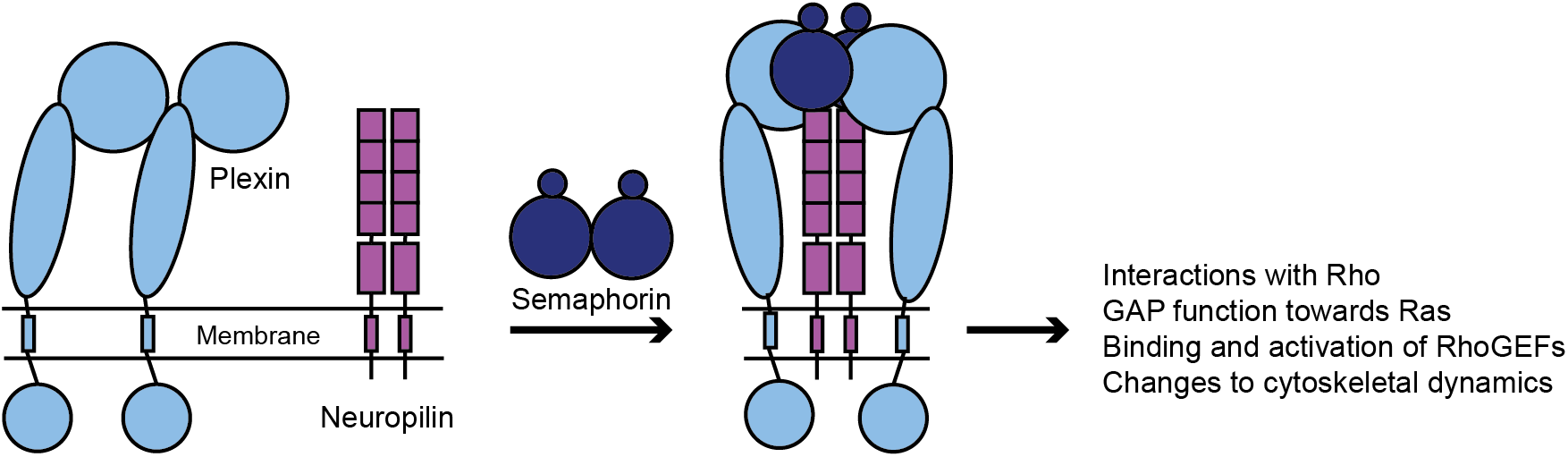
Schematic of hypothesized Plexin-Neuropilin-Class 3 Semaphorin signaling. Using previously available structural and biochemical data, plexins (green) and neuropilins (red) likely form inhibitory homodimers. Addition of a soluble, dimeric class 3 semaphorin induces a tripartite complex formation where neuropilins act as a bridge between plexins and class 3 semaphorins. Conformational changes to the intracellular region of plexins then allow for interactions with GTPases such as Rac1, R-Ras, and Rap1/2 which controls downstream cytoskeletal dynamics.

In this study we probed the interactions of Nrp1, Plexin A2, Plexin A4, and Plexin D1 before and after stimulation with Semaphorin 3A and Semaphorin 3C using PIE-FCCS. Plexin D1 is activated by class 3 semaphorins but has not been investigated with cell biophysical assays. Thus, its oligomer state and potential heterotypic interactions have not been directly assessed. Our results confirm that that in absence of ligand, Nrp1, Plexin A2, and Plexin A4 each form homodimers. In contrast, we discovered that Plexin D1 is monomeric. We also report here for the first time that Plexin A2 and Plexin A4 assemble into a heteromeric complex in the absence of ligand. Intriguingly, each of these homotypic dimer complexes (or lack thereof) was unaffected by Semaphorin 3A or Semaphorin 3C stimulation. A complex between Plexin D1 and Nrp1 was observed following incubation with Semaphorin 3C, however, no interactions were observed following Semaphorin 3A stimulation. The results presented here expand upon previous interaction studies by including multiple receptor and ligand pairs to begin resolving the full interaction profile for this important protein family. Advances in understanding this local network of protein interactions will aid in the development of new therapeutic strategies that target these receptors.

## Methods

### Plasmids and cloning

Each of the full-length human receptor proteins were cloned into pEGFP-N1 and pmCherry-N1 vectors for mammalian expression. Cloning of Plexin A2-eGFP (accession No. O75051) and Plexin A2-mCherry were carried out by inserting the Plexin A2 sequence into EcoR1 and Kpn1 sites of the vectors. The cloning primers are

Forward: 5’ ACTGAATTCATGGAACAGAGGCGGCCCTGGCCCC 3’ and
Reverse: 5’ ACTGGTACCGTGCTCTCAATGGACATGGCAT TAATGAGCTG 3’. Plexin D1-eGFP (accession No. Q9Y4DY) and Plexin D1-mCherry were cloned by using EcoR1 and BamH1 sites in vectors and the Plexin D1 internal Sac1 to amplify two pieces followed by a three-way ligation. The cloning primers are
Forward 1: 5’ AATGAATTCATGGCTCCTCGCGCCGCGGGCGGCGCACCCCTTAGCGCCCGGGCCGCC GCCGCCAGCCCCCCGCCGTTCCAGACGCCGCCGCGGTGCCCGGTGCCGCTGCTGTTG CTGCT 3’;
Reverse 1: 5’ GCACCAGGACCTGGAGCTCGGAGCCTACATGG 3’;
Forward 2: 5’ CCATGTAGGCTCCGAGCTCCAGGTCCTGGTGC 3’;
Reverse 2: 5’ AATGGATCCCGGGCCTCACTGTAGCACTCGTAGATGTTGTCCTCCATCAAAGCCAC 3’.
Cloning of Nrp1-eGFP (accession No. O14786), Nrp1-mCherry, Plexin A4-eGFP (accession No. Q9HCM2), Plexin A4-mCh, Plexin A4ΔSema-eGFP, and Plexin A4ΔSema-mCh was performed as previously described.^34^ The Plexin A4ΔSema mutant deletes residues 39-506 from the full length construct near the N-terminus.

### Cell culture and ligand stimulation

Cos-7 cells were cultured and transiently transfected using standard procedures.^42^ Briefly, culture media consisted of Dulbecco’s modified eagle medium supplemented with 10% fetal bovine serum and 1% penicillin-streptomycin. Cells were passaged at 70-90% confluency to 35 mm glass bottom dishes (Mattek Corporation) for transfection. Approximately 24 hours prior to data collection the cells were transiently transfected with the protein(s) of interest using Lipofectamine 2000 reagent (Thermo Fisher Scientific) and 1.25-5 μg of plasmid DNA. Recombinant human Semaphorin 3C (C636, Bon Opus Biosciences, Milburn, NJ) contains residues 21-738 and is >95% pure. Recombinant human Semaphorin 3A (CX65, Bon Opus Biosciences, Milburn, NJ) contains residues 21-771 and is >95% pure. For stimulation with these ligands, a stock solution (100 μg/mL) was diluted to 500 ng/mL in imaging media and added to receptor-expressing cells approximately ten minutes prior to data acquisition. Data was taken for up to one hour following stimulation.

### PIE-FCCS instrumentation, data collection and analysis

PIE-FCCS data collection was performed as previously described.^8, 43^ Briefly, the custom built set-up uses a 50 ns pulsed continuum white laser source (SuperK Extreme, NKT Photonics, Birkerød, Denmark) split into two wavelengths, 488 nm and 561 nm. These beams are directed through individual optical fibers of different lengths to induce a delay in arrival time relative to each other allowing for PIE and elimination of spectral cross talk between the detectors.^44^ The beam powers were set to 300 nW for 488 nm and 800 nW for 561 nm. The beams were overlapped and directed to the back of the microscope (Eclipse Ti, Nikon Instruments, Tokyo, Japan). These overlapped beams were focused through the objective to a diffraction limited spot on a peripheral membrane area of a Cos-7 cell expressing the eGFP and mCherry labeled receptor constructs. Emitted photons were detected by individual avalanche photodiodes with a 50 μm detection chip (Micro Photon Devices, Bolzano, Italy) and recorded by a time correlated single photon counting module running in time tagged time resolved mode.

For each single cell measurement five acquisitions of ten seconds were recorded at the peripheral membrane area. Each intensity fluctuation was subjected to PIE gating before the auto- and cross-correlation analysis. In an auto-correlation analysis, the intensity at time F(t) was compared to the intensity at a later time F(t+τ) and the self-similarity as a function of the later time allowed for interpretation of quantitative information such as diffusion and the number of particles. Intensity fluctuations were separated into 10 μs bins and subjected to the correlation algorithm in Eq. 1, which normalizes the intensity change to the square of the average intensity.^45–47^ Cross-correlation uses the intensity fluctuations which occur simultaneously in both channels to infer interaction of species. Here, the correlation algorithm is represented by Eq. 2 and the ratio of the cross-correlation amplitude to the auto-correlation amplitude indicates the proteins in complex, limited by the lower population molecule.^45^

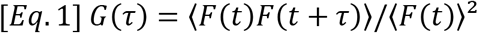

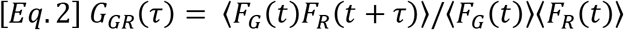

The five acquisitions from each single cell measurement were averaged together to remove perturbations such as cell movement or bright clusters. Once individual curves from each cell are averaged, a least squares fitting to a 2D diffusion model is used, and includes fitting parameters for the triplet state, Eq. 3.

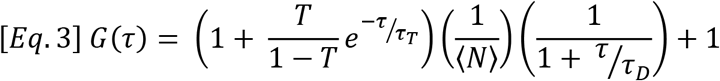

With auto-correlation it is possible to infer protein diffusion by using the timing of intensity fluctuations. The half value decay time (lag time, τD) can be used in Eq. 4 to determine the effective diffusion coefficient.^46^ Diffusion will be affected by protein molecular weight and interactions with other molecules.

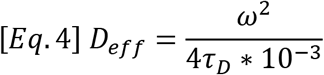

With the addition of a second detection channel the co-diffusion of two proteins can be analyzed by PIE-FCCS. The overlapping laser beams create a defined area for both eGFP and mCherry tagged proteins and their intensity fluctuations will occur simultaneously as they pass through the illuminated area.^45–48^ Following cross-correlation by the algorithm stated above, the amplitudes can be compared as shown in Eq. 5.^45^

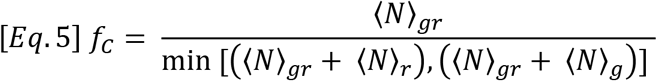

An ideal system would have a fraction correlated (*f*_c_) of zero for a non-interacting species and a fraction correlated of one for an interacting species. However, we must take certain considerations into account when interpreting live cell fluctuation results. Our lab’s previous publications show a set of control constructs with a myristoylation anchor and repeated FKBP domains fused to a fluorescent protein allowing for interpretation of *f*_c_ values for monomers, dimers, and higher order oligomers.^8, 49^ For homotypic interactions, *f*_c_ values below 0.09 indicate monomeric species, 0.09 < *f*_c_ < 0.17 indicate dimeric species, and those above 0.17 indicate higher order oligomers.

### Western blotting

Samples for Nrp1, Plexin A2, Plexin A4, and Plexin D1 endogenous expression in Cos-7 were collected 24 hours after passaging. Samples for transient expression of Nrp1-eGFP, Plexin A2-eGFP, Plexin A4-eGFP, and Plexin D1-eGFP in Cos-7 were transfected with 2.5 μg plasmid DNA 24 hours after passaging and collected 24 hours post-transfection. Cells were lysed using RIPA lysis buffer supplemented with protease inhibitors (benzamidine, leupeptin, and PMSF). Western blotting experiments with these samples was carried out to confirm the expression of the samples. Primary antibodies used are Nrp1 (Cell Signaling, Cat& 3725), Plexin A2 (R&D, Cat& MAB5486), Plexin D1 (R&D, Cat& AF4160), Plexin A4 (R&D, Cat& MAB5856) and FLAG (Sigma-Aldrich, Cat& F1804).

## Results

### PIE-FCCS shows that Nrp1 forms multimers, Plexin A2, and Plexin A4 form homodimers, Plexin D1 does not self-associate

To measure the spatial organization of Nrp1, Plexin A2, Plexin A4, and Plexin D1 in cells, we first expressed them individually by co-transfection of the eGFP and mCherry fusion constructs to determine their degree of homodimerization. PIE-FCCS data was collected from single live Cos-7 cells expressing the tagged protein of interest at surface densities ranging from 85-1245 molecules/μm^2^. From each cell measurement we quantified expression of eGFP- and mCherry-labeled protein, the 2D mobility of the receptors in the plasma membrane, as well as the degree of association using the fraction of cross-correlation, *f_c_*.^8^ In order to ensure endogenous receptors would not interfere with the correlation analysis we used Western blotting to confirm that transiently transfected plasmids are expressed at increased levels compared to endogenous expression (Figure S1). PIE-FCCS directly quantifies the expression level of the FP fusion in each single cell measurement, which varied between 85 and 1245 molecules/μm^2^.

The *f*_c_ values were used to determine the degree of oligomerization for each receptor (Figure 2A). The Nrp1 data had a median of 0.14, which is consistent with strong dimerization, however, the wide distribution of *f*_c_ values over 0.20 suggests that Nrp1 can also form small homotypic multimers as reported previously.^36–37^ The median *f*_c_ value of 0.14 for Plexin A4 is consistent with dimerization, as reported previously by PIE-FCCS.^34^ The Plexin A2 cross-correlation had a lower median (*f*_c_ = 0.09) indicating a weaker dimer affinity compared to Plexin A4. To the best of our knowledge, there has been no investigation of the oligomerization state of Plexin D1, except for a computational prediction that the isolated transmembrane helix is expected to dimerize to a similar extent as other plexin transmembrane domains.^50^ The near zero fraction correlated observed here (median *f*_c_ = 0.01) suggests that full length Plexin D1 does not dimerize in the given concentration range. The diffusion coefficients for each receptor support the interpretations of the *f*_c_ values (Figure 2B), with higher mobility observed for monomeric Plexin D1 compared to Plexin A2 and Plexin A4. Nrp1 has an average diffusion coefficient of 0.26 μm²/sec, consistent with the formation of multimers as this is significantly slower than dimeric Plexin A2 and Plexin A4, where the average diffusion coefficients are 0.39 and 0.37 μm²/sec, respectively. Plexin D1 has the fastest average diffusion coefficient, 0.59 μm²/sec, adding to the evidence that it is monomeric.

**Figure 2.**
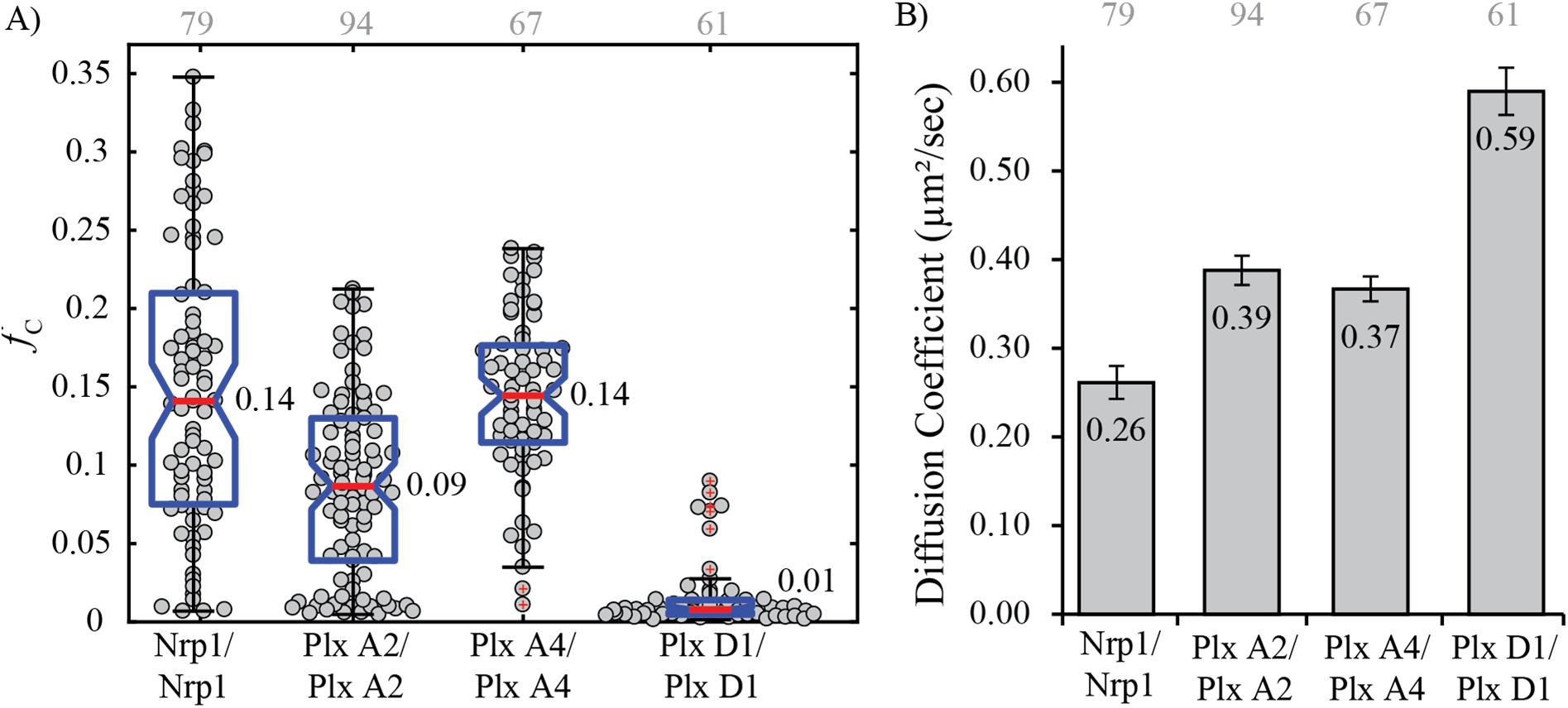
Homotypic interaction of Nrp1, Plexin A2, Plexin A4 and Plexin D1. A) Fraction correlated for Nrp1, Plexin A2, and Plexin A4 fall in the range of homodimers, while Plexin D1 diffuses as a monomer. Grey numbers above each column represent the number of single cells analyzed. B) The average diffusion coefficients agree with the cross-correlation results, where Plexin D1 (monomer) diffuses at a faster rate than the dimers, but the slow diffusion for Nrp1 suggests multimers may form as well, possibly involving interactions other endogenous proteins.

### Nrp1 does not interact significantly with plexins in the absence of semaphorin ligand

Nrp1 is involved in class 3 semaphorin signaling as well as other ligands like VEGF but cannot transduce the signal without expression of additional receptors (i.e. plexins, VEGFR, MET). However, conflicting reports exist regarding the interactions of these receptors in heteromeric complexes before and after stimulation. Man *et al.*^26^ presented data suggesting the interaction of Plexin A2, Plexin D1, and Nrp1 following stimulation with Semaphorin 3C while others observed different heteromeric complexes prior to stimulation or even no interactions at all.^20, 28, 30–31^ To measure the heterotypic interactions between Nrp1 and the plexin receptors, each plexin-eGFP construct was co-expressed with Nrp1-mCh. Single cell PIE-FCCS data was collected for each combination to determine the degree of association and 2D mobility. The *f*_c_ values for Nrp1 co-expressed with each plexin construct each had a median value of 0.01 (Figure 3A). This lack of cross-correlation indicates that unstimulated receptors have negligible propensity to dimerize with Nrp1 in the live cell plasma membrane. The average diffusion coefficient of Nrp1 expressed with Plexin D1 showed a modest increase from 0.26 to 0.34 μm²/sec compared to when it was expressed without Plexin D1 (Figure 3B). This suggests that Nrp1 may form oligomers when expressed alone, but shifts toward dimerization when in the presence of co-receptors as previously reported.^36–37^ The diffusion of the plexin receptors is not drastically altered in the presence of Nrp1 except in the case of Plexin A4 which has a significant decrease in average diffusion coefficient, 0.37 to 0.31 μm²/sec (Figure S2). It is possible that Plexin A4 interacts weakly with Nrp1 oligomers before stimulation, as evidenced by the comparatively large distribution of *f*_c_ values (Figure 3A). With Nrp1 acting as a co-receptor for many other receptors (e.g.VEGFR2), the lack of cross-correlation in our assay may also be due to a competition between plexins and other endogenous receptors, i.e. the plexin binding to Nrp1 is too weak of off-compete these interactions.

**Figure 3.**
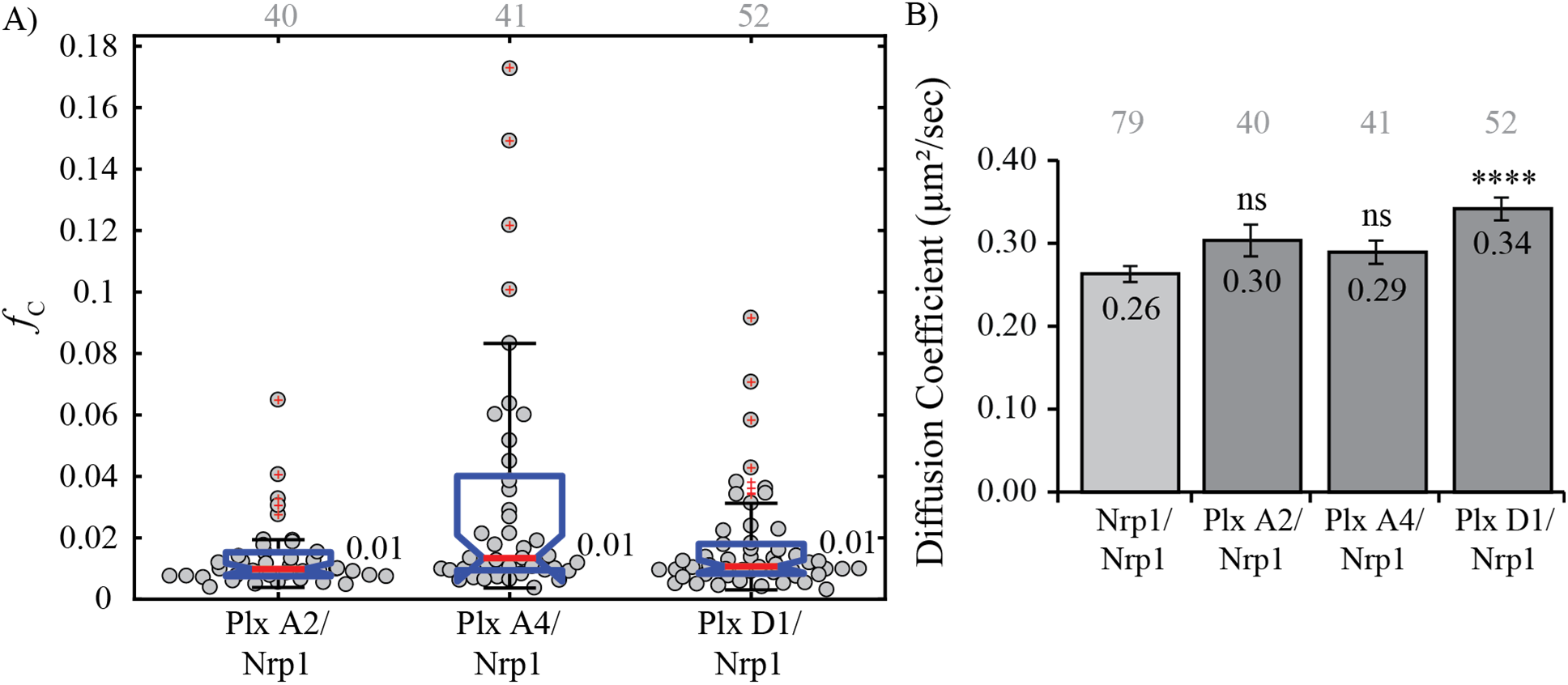
Heterotypic interaction of Nrp1 with plexin receptors. A) The fraction correlated indicates no interaction between any receptor combinations under non-stimulatory conditions. Grey numbers above each column represent the number of single cells analyzed. Data marked with red + are regarded as outliers and are not included in the analysis. B) Comparison of effective diffusion coefficient of Nrp1 when expressed alone (light grey) or co-expressed (dark grey). Comparison with Nrp1-mCh diffusion in the homodimer experiments shows that Nrp1-mCh diffusion is significantly increased when co-expressed with Plexin D1 (p<0.0001), but not with Plexin A2 or Plexin A4.

### Class A Plexins can form heterodimers via their sema domain, suggesting a heterotypic interaction model

Most studies on plexin/neuropilin/semaphorin signaling have focused on one receptor-ligand pair or the interaction with Nrp1. Therefore, little information has been reported for the heterotypic interactions of the plexins themselves. The previous report by Man *et al.*^26^ suggested such interactions in glioblastoma multiform samples. However, due to the endogenous expression of Semaphorin 3C no unstimulated data was obtained. In a 2003 report, Plexin A1 and Plexin B1 were suggested to associate via their cytoplasmic domains.^51^ In another study, Smolkin *et al.*^29^ determined that Plexin A4 and Plexin D1 were not associated when unstimulated, but could form a complex when in the presence of Semaphorin 3C.

To determine the degree of interaction between each plexin receptor pair we conducted pair-wise co-expression of each receptor combination in Cos-7 cells and collected PIE-FCCS data. The median *f*_c_ for Plexin D1 co-expressed with either class A plexin is approximately zero (Figure 4A). This indicates that neither class A plexin forms a complex with Plexin D1 under non-stimulatory conditions. However, the median *f*_c_ value for Plexin A2 and Plexin A4 is 0.08, suggesting the presence of heterodimers in live cells. The average diffusion coefficient of both class A plexins is unchanged when expressed alone or with the other class A plexin (Figure 4B). This allows us to conclude that the complex formed is most likely a heterodimer and not a larger multimer. This is the first observation, to the best of our knowledge, that class A plexins can be involved in heterodimers with other class A plexins prior to ligand stimulation.

**Figure 4.**
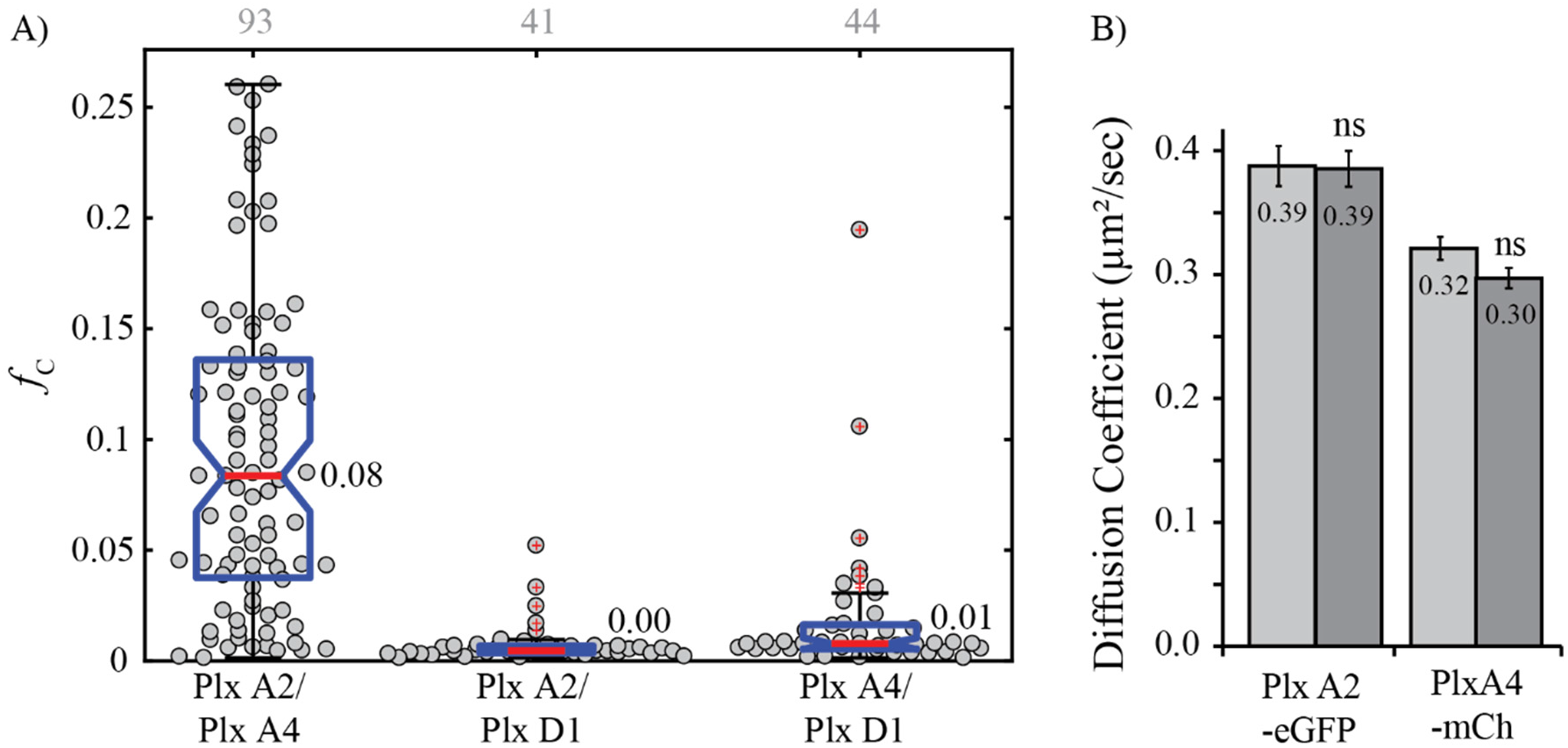
Heterotypic interactions of plexins. A) Fraction correlated for each combination indicates that Plexin A2 and Plexin A4 form a hetero-dimer and neither interacts with Plexin D1. Grey numbers above each column represent the number of single cells analyzed. B) Diffusion coefficient change for Plexin A2 and Plexin A4 when expressed alone (light grey) or together (dark grey) likely indicating that neither forms a complex larger than a homo- or heterodimer.

Based on a crystal structure from Kong *et al,*^35^ class A plexins form the inhibitory homodimer in a “head to stalk” fashion, with the sema domain acting as the “head” and the PSI2/IPT2 domain as the “stalk” (cartoon shown in Figure 1). We co-expressed a mutant Plexin A4 with a complete deletion of the sema domain (Plexin A4ΔSema), previously used by Marita *et al.*^34^ to determine if the heterodimer was also dependent on the sema domain. Using this construct in combination with WT Plexin A2 shows a dramatic increase in the median *f*_c_ value, from 0.08 to 0.23 (Figure S3A). The reason for the dramatic increase in cross-correlation is likely due to the reduced competition with the Plexin A4 homodimers and possibly the release from an autoinhibited structure (see discussion). When Plexin A4 lacks the sema domain it can no longer form a homodimer, leading to the combinations A2:A2, A2:A4ΔSema, and an A4ΔSema monomer (Figure S3B). The Plexin A4 homodimer is now abrogated, reducing competition with the heterodimer, and causing the dramatic increase in co-diffusion. These experiments show the necessity for heterodimerization analysis by PIE-FCCS to begin building a model that incorporates the full complexity of membrane protein interaction networks.

### Semaphorin 3C induces complex formation for Plexin D1 and Nrp1, while Semaphorin 3A does not induce a detectable interaction

Few data have been reported on whether the homotypic interaction of plexins is changed following ligand stimulation by Semaphorin 3C or Semaphorin 3A, except that direct binding to Nrp1 can occur.^20, 31^ Following on the approach in Man *et al.^26^*, which delivers exogenous Semaphorin 3C to cells in a dose dependent manner, we incubated Cos-7 cells expressing individual receptors with 500 ng/ml of recombinant human Semaphorin 3C or Semaphorin 3A. Incubation times for previous experiments varied from minutes^29^ to days^26^ depending on the context of the experiment. Because we were interested in the early events of receptor interaction at the membrane rather than downstream signaling events, we collected PIE-FCCS data between 10 and 70 minutes after ligand stimulation. The average *f*_c_ value for each receptor was unchanged following stimulation indicating that homotypic oligomerization was not significantly enhanced or disrupted (Figure 5). However, both Plexin A4 and Plexin D1 showed a significant increase in average diffusion coefficient following ligand stimulation, with Semaphorin 3A (0.37 to 0.44 μm²/sec) and Semaphorin 3C (0.59 to 0.68 μm²/sec), respectively (Figure S4). These diffusion changes may be interpreted as a change in conformation and/or unbinding of an endogenous (unlabeled) protein, but it is unlikely that the homo-oligomerization state is affected by stimulation. This result is consistent with the crystal structure of the Plexin A2-Nrp1-Semaphorin 3A complex^35^ where the interactions of Nrp1 are predominantly with the Plexin A2-bridging semaphorin, with few or any Nrp1 domain contacts with Plexin.

**Figure 5.**
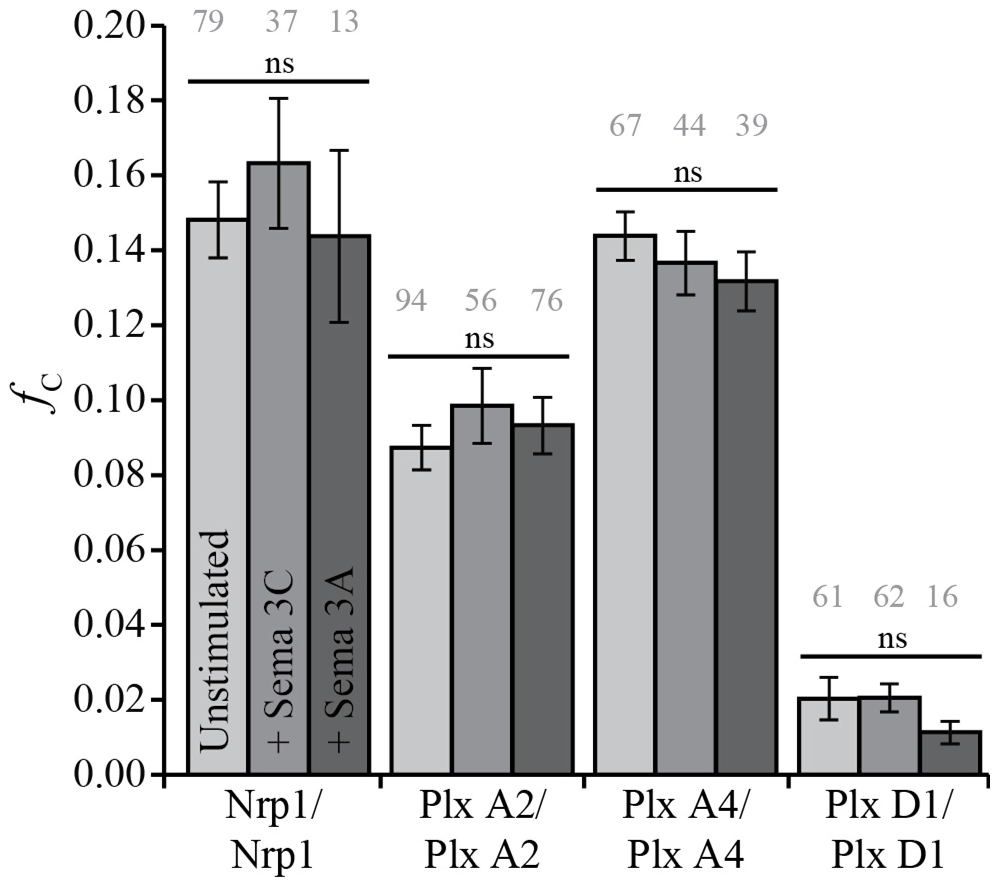
Homotypic interaction of Nrp1, Plexin A2, Plexin A4, and Plexin D1 following stimulation with Semaphorin 3C and Semaphorin 3A. Cells expressing homotypic receptor combinations were incubated with 500 ng/ml of Semaphorin 3C or Semaphorin 3A 10 minutes prior to data acquisition. Fraction correlated is unchanged from non-stimulatory conditions. Grey numbers above each column represent the number of single cells analyzed.

We next tested the interaction between Nrp1 and each plexin receptor in the presence of semaphorin ligands. Each receptor combination was co-expressed in Cos-7 cells and PIE-FCCS data was collected as in the previous experiments. Using the same concentration and incubation conditions from the previous section, the co-expressed receptors were stimulated with recombinant Semaphorin 3C. The *f*_c_ values for Nrp1-mCh co-expressed with each plexin-eGFP construct are reported in Figure 6A. Plexin A2-Nrp1 and Plexin A4-Nrp1 each have a median *f*_c_ value of 0.01, which is similar to the unstimulated values, indicating a lack of interaction. Both Man *et al.^26^* and Toyofuku *et al.^32^* observed Plexin A2/Nrp1 Co-IP following Semaphorin 3C stimulation, but analysis of PIE-FCCS data shows no interaction at this ligand concentration and receptor expression range (85-1245 molecules/μm²) in the live cell plasma membrane. Following Semaphorin 3C stimulation, Plexin D1-Nrp1 did show substantial increase in heterodimerization (*f*_c_ = 0.13). These changes in oligomerization state of Plexin D1 and Nrp1 were supported by changes in the effective diffusion coefficient (Figure 6B and C). The average diffusion coefficient of Plexin D1 decreased by 20% (0.62 to 0.50 μm²/sec), indicating increased molecular weight and a shift from monomer to heteromeric complex (Figure 6B). The Nrp1 diffusion coefficient was significantly higher than in the homodimer experiments (0.31 compared to 0.26 μm²/sec) but does not significantly decrease upon stimulation with Semaphorin 3C (0.34 to 0.31 μm²/sec). This result is consistent with a shift from Nrp1 homomultimers to heteromeric complexes (Figure 6C). Figure S5 shows that the *f*_c_ distribution for each combination of plexin receptors was relatively unchanged following Semaphorin 3C stimulation.

**Figure 6.**
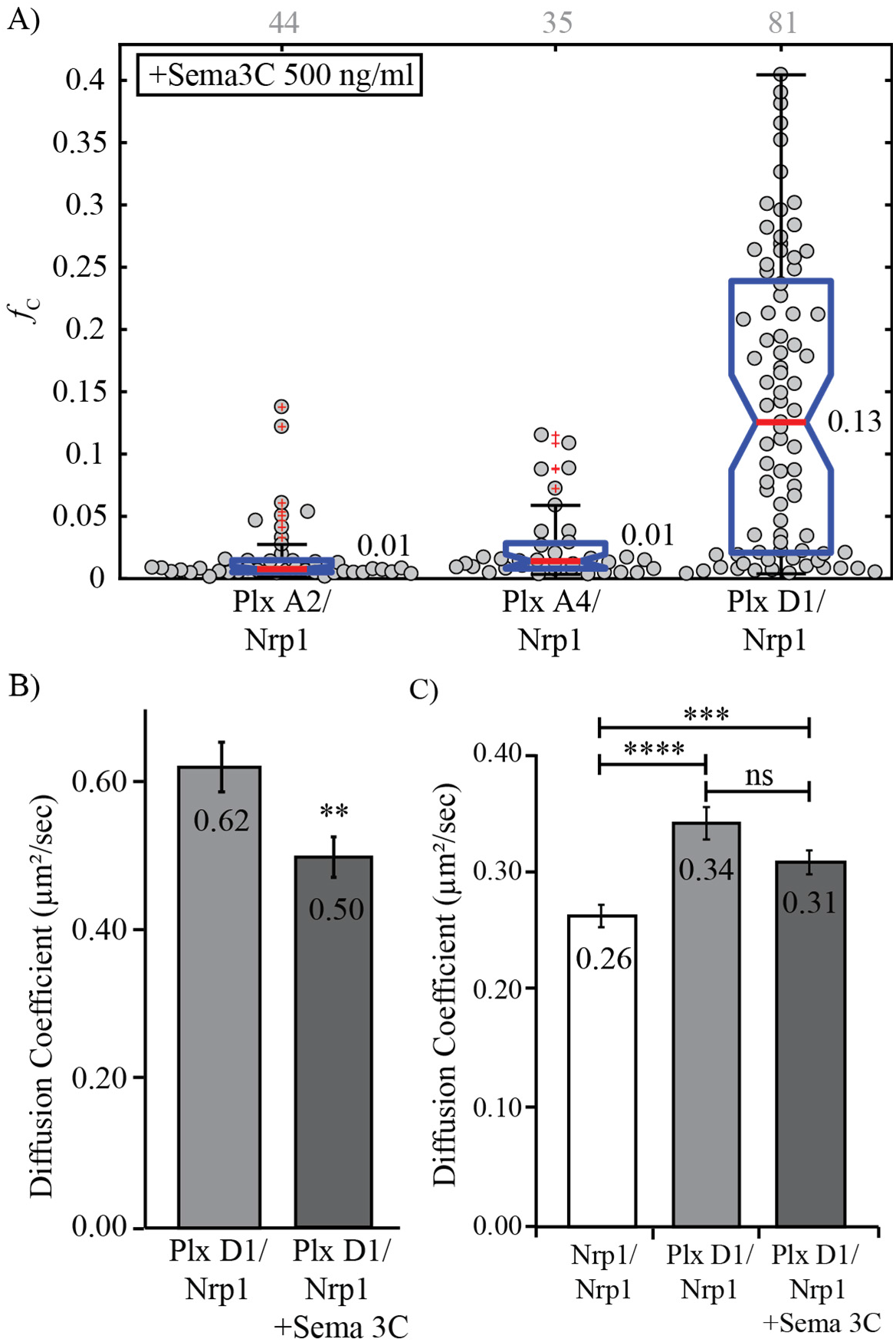
Heterotypic interaction of Nrp1, Plexin A2, Plexin A4, and Plexin D1 following stimulation with Semaphorin 3C. A) Fraction correlated for Nrp1 co-expressed with each plexin receptor. Plexin D1 and Nrp1 exhibit extensive dimerization. Grey numbers above each column represent the number of single cells analyzed. B) Diffusion change for Plexin D1-eGFP when co-expressed with Nrp1. Stimulation with Semaphorin 3C significantly decreased (p<0.01) the average diffusion coefficient indicating increased molecular weight and oligomer state. C) Diffusion change for Nrp1-mCh alone or when co-expressed with Plexin D1. Again, the average diffusion coefficient is significantly increased from expression alone, but not significantly decreased from the unstimulated co-expression.

Various experiments have suggested that heteromeric complexes of Nrp1 and class A plexins form after stimulation.^10, 12, 31, 38–41, 52–55^ As demonstrated above, before stimulation neither Plexin A2 nor Plexin A4 were observed to form a heteromeric complex with Nrp1. Using the same conditions as above, Cos-7 cells co-expressing each combination of Nrp1 and plexin receptor were stimulated with Semaphorin 3A then probed with PIE-FCCS measurements to assess any changes in mobility and association. Figure 7A reports the *f*_c_ values for each set of receptors. No drastic changes in co-diffusion were observed for any combination. In Figure 7B we have compared the average fraction correlated for these combinations before and after stimulation and observed small but statistically significant increase for Plexin A2-Nrp1 and Plexin A4-Nrp1 when exposed to the ligand (0.01 to 0.05, 0.03 to 0.07, respectively). Average diffusion coefficients for Plexin D1 and Nrp1 were unchanged compared to homotypic and heterotypic interaction rates, however, Plexin A4 and Plexin A2 diffusion coefficients decreased (Figure 7C). Both Plexin A2-eGFP and Plexin A4-eGFP have their lowest average diffusion coefficient when expressed with Nrp1 and stimulated with Semaphorin 3A compared to expression of the plexin alone, 0.39 to 0.30 μm²/sec and 0.37 to 0.28 μm²/sec, respectively. Nrp1-mCh diffusion was not significantly changed under any condition. Overall, these data do not give any clear indication of a transition to heterodimer after Semaphorin 3A binding as was seen for Plexin D1 and Nrp1 following Semaphorin 3C binding. However, some caution is advised when interpreting these negative results. PIE-FCCS measurements of heterodimers is affected by the stability and dynamics of the heterodimer as well as any competition with homodimers and heterotypic interactions with other endogenous receptors. Until the full network of membrane protein interactions can be resolved, it is difficult to rule out low affinity interactions based on negative PIE-FCCS studies. Alternatively, it is also possible that three receptors may be necessary to form a stable, signaling complex and that pairwise expression of exogenous receptors is insufficient to drive the formation of the full signaling complex. This type of heteromeric complex has been suggested in previous studies, but not observed directly in live cell biophysical assays.^10, 54, 56–57^ Future work using three-color labeling could help resolve these putative assemblies.

**Figure 7.**
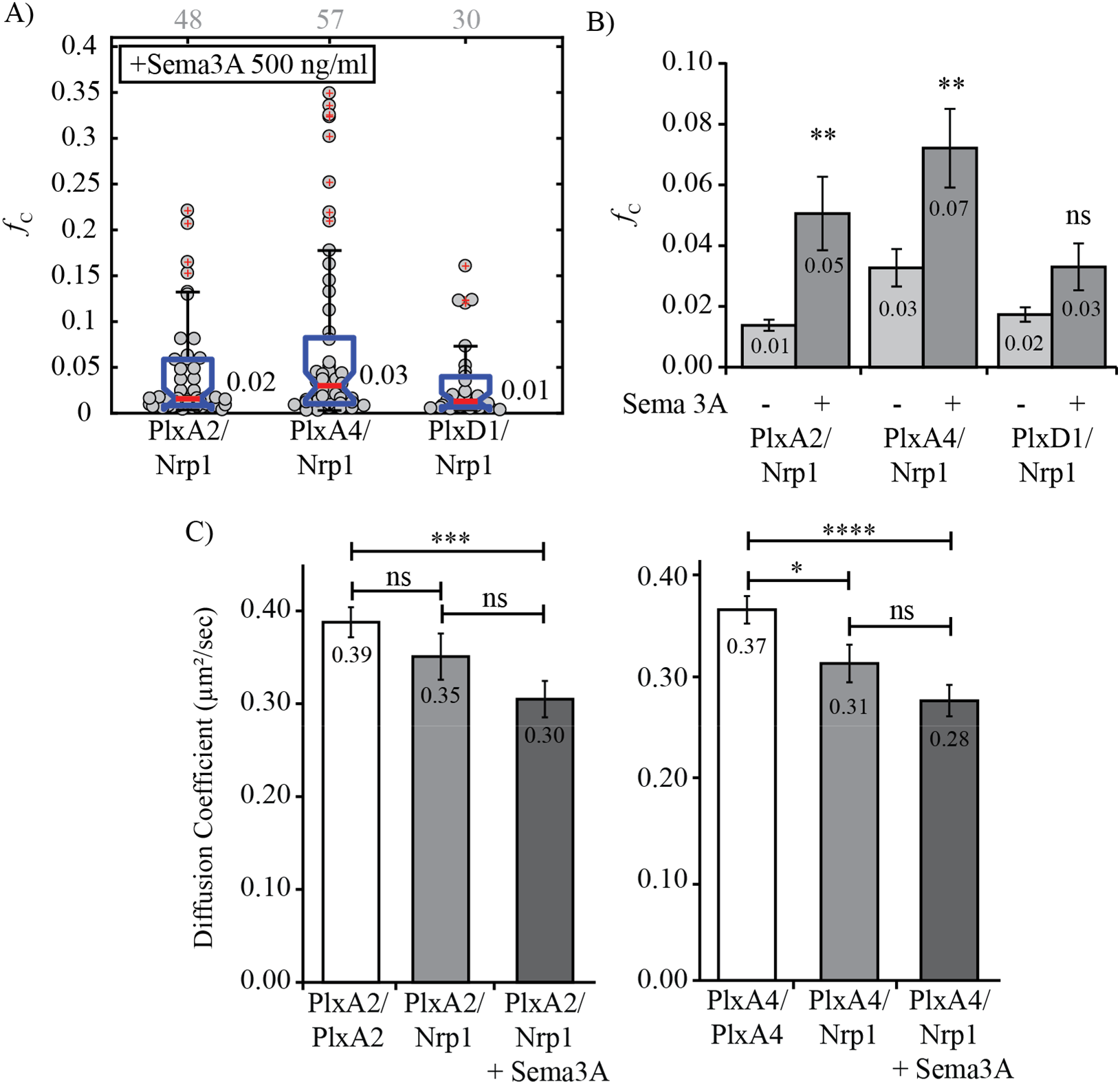
Heterotypic interaction following Semaphorin 3A stimulation. A) Fraction correlated after stimulation. No apparent increase is observed like that of Plexin D1-Nrp1 following Semaphorin 3C stimulation. Grey numbers above each column represent the number of single cells analyzed. B) Changes in average fc value following stimulation. Plexin A2-Nrp1 and Plexin A4-Nrp1 have significant increases in correlation while the Plexin D1-Nrp1 interaction is unchanged. C) Diffusion change for Plexin A4-eGFP and Plexin A2-eGFP when co-expressed with Nrp1 and stimulated with Semaphorin 3A. Both Plexin A4 and Plexin A2 have significantly decreased average diffusion coefficients adding to evidence that a weak/transient interaction is formed.

## Discussion

Semaphorin 3A and Semaphorin 3C are needed for normal development, but disruption after the embryonic developmental stages can lead to various disease states. Depending on the tissue type, Semaphorin 3A stimulation can enhance or inhibit angiogenesis and migratory pathways in tumor cell populations.^6, 58–59^ Multiple studies have shown that reduction of Semaphorin 3A expression occurs in later stages of cancers (including breast and prostate) and that exogenous Semaphorin 3A leads to reduced metastasis and angiogenesis.^27, 60–65^ Therefore, it is an important ligand to study as a potential anti-migratory therapeutic factor.^66^ During development, Semaphorin 3C downregulation in cardiac tissue is related to certain types of congenital heart disease.^21^ In later stages of life, Semaphorin 3C overexpression is involved in multiple cancer types, including glioblastoma,^26^ lung,^67^ gastric,^68^ ovarian^69^ and prostate.^27–28, 65^ Angiogenesis can also be increased in the presence of semaphorins, and receptors for Semaphorin 3C, particularly Plexin D1, are upregulated in the tumor vasculature making it a potential drug candidate.^30, 70^ In order to fully understand the effects of these ligands, we must elucidate whether and how their receptors interact before stimulation and how their configurations are altered upon ligand-receptor complex formation. Previous work suggested that Semaphorin 3C, Plexin A2, Plexin D1, and Nrp1 form a complex in glioma stem cells,^26^ while numerous studies have indicated the interaction of Semaphorin 3A, plexins, and Nrp1. Our goal here was to determine the possible interaction modes in a live cell environment. Using PIE-FCCS, we were able to analyze these homotypic and heterotypic interactions of membrane receptors before and after ligand stimulation.

Our work extends previous PIE-FCCS studies of Plexin A4 dimerization to a larger set of receptors and ligands for which co-existing homodimers and heterodimers could compete for binding.

We first confirmed that Nrp1, Plexin A2, and Plexin A4, all form homodimers in the absence of ligand stimulation as previously reported.^34–37^ We next determined that the full-length Plexin D1 protein is a monomer, which to the best of our knowledge, is reported here for the first time. However, the homodimerization of Plexin D1 was inferred based on computational prediction of reasonably strong interactions between the transmembrane helical domains.^50^ Different configurational states have been presented by crystallography and cryo-EM for the extracellular region of plexins over the last several years,^35, 39^ and the functional autoinhibition of such states can be relieved by truncation of the extracellular domains. For example, deletion of the Plexin A1 sema domain converts the protein from an autoinhibited form to a constitutively active protein (in absence of ligand).^55^ In principle, it is possible that Plexin D1 may undergo an inactive to active state transition without the need for homodimerization.^7, 71–72^

The receptors examined here do not form heterotypic interactions in the absence of ligand, except for Plexin A2 and Plexin A4. The class A plexins have conserved residues which may contribute to heterodimerization; however, these interactions had not been reported prior to the present study. Deletion of the sema domain from Plexin A4 (Plexin A4ΔSema) inhibits the homodimerization as we previously reported.^34^ When Plexin A4ΔSema was co-expressed with WT Plexin A2 there was a dramatic increase in the amount of cross-correlation and thus the degree of heterodimerization. This effect is ascribed to the fact that there was no longer competition from Plexin A4 homodimers, allowing for a greater number of monomeric Plexin A4ΔSema molecules to form A2:A4 heterodimers. The strong interaction between Plexin A2 and Plexin A4ΔSema also suggests the dimerization is between the Sema domain of A2 and the “stalk” region of Plexin A4ΔSema. This is consistent with the recent cryo-EM structures of the Plexin ectodomains.^35^ These results support a model of heterotypic interactions where multiple binding partners and affinities must be taken into account to fully understand signaling. In addition, interactions such as these must be considered when disrupting or mutating receptors for disease related research as signaling may still occur through related endogenous proteins.

After establishing the ligand-independent interactions, we now discuss the receptor interactions following stimulation with semaphorin ligands. PIE-FCCS shows Semaphorin 3C stimulation influences the interaction of Plexin D1 and Nrp1, which was only indirectly observed in previous studies.^20, 30^ Our findings indicate that Semaphorin 3C signal transduction may not utilize Plexin A2 as a receptor, even though it appears to form a complex when observed by co-IP or AP-binding assay.^26, 32^ Figure 8 shows a model of the Plexin D1-Nrp1-Semaphorin 3C interaction. Semaphorins are inherent dimers which have been shown to bind their receptors in a 2:2 stoichiometry as suggested by crystal structures.^39, 73–74^ Taking this and our PIE-FCCS analysis into account there are various interactions which may occur following Semaphorin 3C stimulation. The first option is a 1:1:2 Plexin D1-Nrp1-Semaphorin 3C complex. Here the median *f*_c_ value falls within the range expected for simpledimerization,^49^ but the values may be altered by monomeric Plexin D1 and dimeric Nrp1. If the Nrp1 homodimer has a high binding affinity it is also possible that Semaphorin 3C causes a 1:2:2 complex where a monomeric Plexin D1 binds to a Nrp1 dimer upon stimulation (Figure 8). In addition, Plexin A2-Nrp1-Semaphorin 3A form a 2:2:2 complex in the low/medium resolution crystal structure and this receptor-ligand complex may have the same stoichiometry as shown in Figure 1.^35^ Importantly, this Nrp1 domain only makes substantial contacts with the dimeric Semaphorin 3A and not with the Plexin A2 sema domain. Although the resolution of the complex structure was medium/low at 7 Å and Nrp1 domains a2, b1, and b2 were not seen in the crystal, the lack of Nrp1-Plexin A2 interactions in the PIE-FCCS data is consistent with the negligible effect of ligand binding on Plexin A2 and Plexin A4 homodimerization. In the 2:2:2 crystallographic structure there were no direct interactions between the Plexin A2 sema domains. This is consistent with a model in which the sema-PSI2/IPT2 domain interactions between plexins are replaced by sema domain interactions between plexin and semaphorin in the complex. Future experiments will need to be performed to determine the stoichiometry of receptors within the signaling complex as well as the time scales of the formation and disruption of the complex.

**Figure 8.**
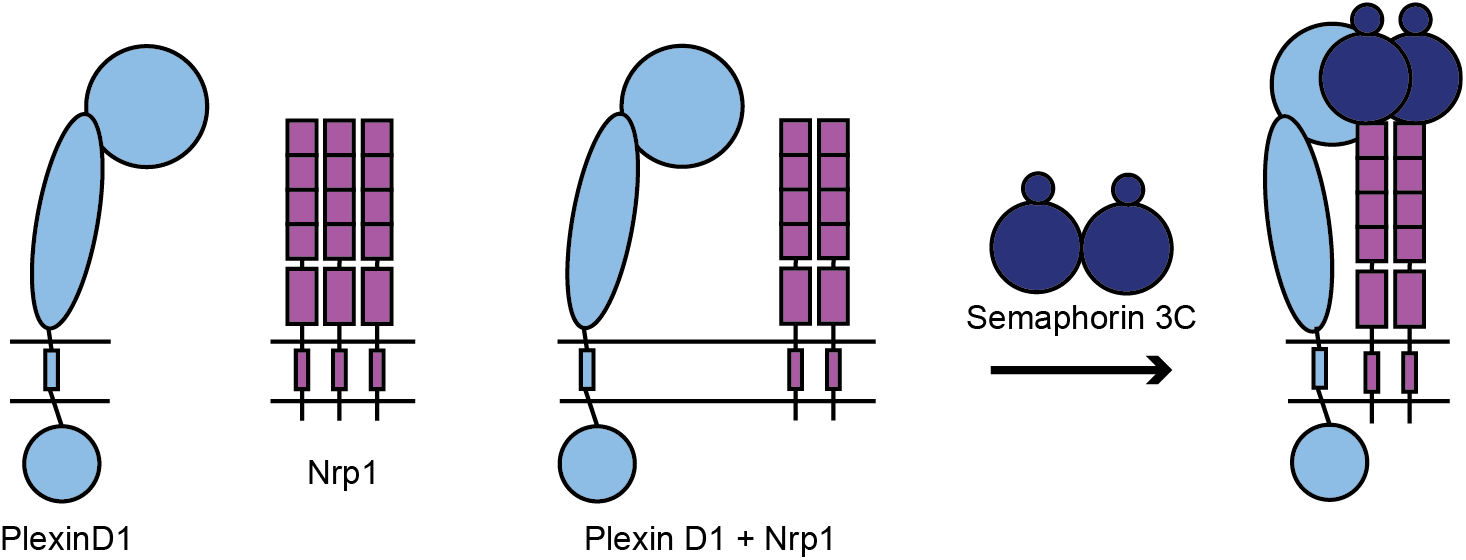
Possible stoichiometry of Plexin D1-Nrp1-Semaphorin 3C complex. Plexin D1 diffuses as a monomer while initially Nrp1 diffuses as a dimer or multimer. Upon co-expression Nrp1 likely shifts toward dimers. Due to Plexin D1 diffusing as a monomer it is possible to form a 1:2:2 complex (right) using a Plexin D1 monomer to induce signaling rather than a dimer seen for Class A plexins.

While previous reports have suggested that Nrp1 and class A plexins form a complex after stimulation with Semaphorin 3A, our analysis by PIE-FCCS does not provide strong supporting evidence. A large increase in the *fc* distribution, like that observed for Plexin D1-Nrp1-Semaphorin 3C, was not observed for Nrp1 and Plexin A2 and A4 receptors when incubated with Semaphorin 3A. There was a small but statistically significant increase in the mean *f*_c_ value for Plexin A2-Nrp1 and Plexin A4-Nrp1 with Semaphorin 3A as well as a decrease in receptor mobility as seen in the diffusion coefficients (Figure 7B-C). There are several possible explanations for these results. First, the timescale of the association could be short. Our single cell measurements were performed during a period of 10-60 minutes after ligand addition. It could be that the formation of the complex is transient and thus appears weak in the average cross-correlation measurements. A second reason could be that there are multiple competing interactions with endogenous proteins. Due to the relative affinities of these competing interaction partners, there may be an ideal set of expression levels under which the heteromeric complex reported in the co-IP and crystallography studies is visible via PIE-FCCS. As stated above, PIE-FCCS measurements of heterodimerization are affected by the stability and dynamics of the heterodimer as well as any competition with homodimers and heterotypic interactions with other endogenous receptors. Until the full network of membrane protein interactions can be resolved, it is difficult to rule out low affinity interactions based on these negative PIE-FCCS results.

Finally, we report here that Plexin A2 and Plexin A4 form heterodimers, and that the extent of heterodimerization is unaffected by ligand binding. The observed A2/A4 heterodimer suggests that ligand binding induces a conformational change that activates the protein rather than driving dimerization per se. Studies of Plexin B1 have led to a model in which pre-existing dimers are not just conformationally altered upon ligand binding, but that plexin-semaphorin 2:2 heterodimers may also associate to form larger order complexes.^39^ Our work suggests the possibility of a fundamental difference in the activation mechanism of plexin A, B and D subfamilies.

Overall, the work here has investigated only a subset of the potential interactions within the plexin/neuropilin/semaphorin protein family. More receptors and ligand combinations will need to be analyzed to establish a more expansive and holistic understanding of how membrane protein-protein interactions regulate plexin/semaphorin signaling. Due to the large number of ligands and receptors (seven class 3 semaphorins, nine plexin receptors, and two neuropilins) this is a time- and resource-intensive undertaking. The work we presented here lays the groundwork for such a comprehensive study. PIE-FCCS is an ideal method for quantifying these interactions in a live cell environment. Combined with cell signaling and high-resolution structure studies, it will be possible to resolve the function role of receptor homo- and heterodimerization in this important signaling axis. This work will also reveal how dysregulated signaling by plexins and neuropilins influence disease states, which will enable new approaches for designing therapeutic strategies.

## Acknowledgements

We thank Dr. ZhenLu Li for helpful discussions during the course of these studies. This work was supported with funding from the National Institutes of Health under grant number R01EY029169 (AWS and MB), the National Science Foundation under grant number CHE-1753060 (AWS). This work was also supported by the NIH/NINDS R01NS094199 and R01NS092641, VeloSano, Amy Post Foundation, Case Comprehensive Cancer Center, and the Cleveland Clinic (JSY).

