## Supplemental Information for "Resolving the Interactions between Class 3 Semaphorin Receptors in Live Cells"

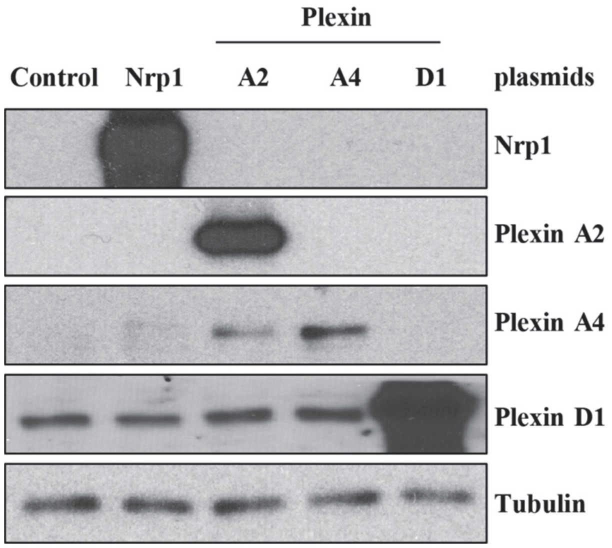


***Figure S1.*** *Schematic of hypothesized Plexin-Neuropilin-Class 3 Semaphorin signaling. Using previously available structural and biochemical data, plexins (green) and neuropilins (red) likely form inhibFigure S1. Western blot of endogenous and transfected receptors. In each case of single transfection, transient receptors are expressed at higher levels than any endogenously present proteins. A band for transient Plexin A2 is present in the Plexin A4 transfected cells most likely due to non-specificity of the antibody.*

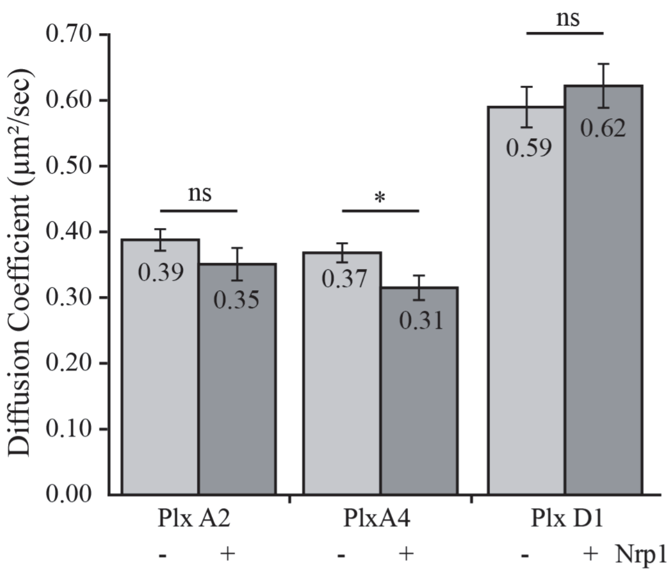


***Figure S2.*** *Diffusion coefficients of plexins alone or co-expressed with Nrp1. Pairwise comparison of plexin-eGFP with and without co-expression of Nrp1 shows that only Plexin A4 has a significant decrease ($p<0.05$) in diffusion in the presence of Nrp1.*

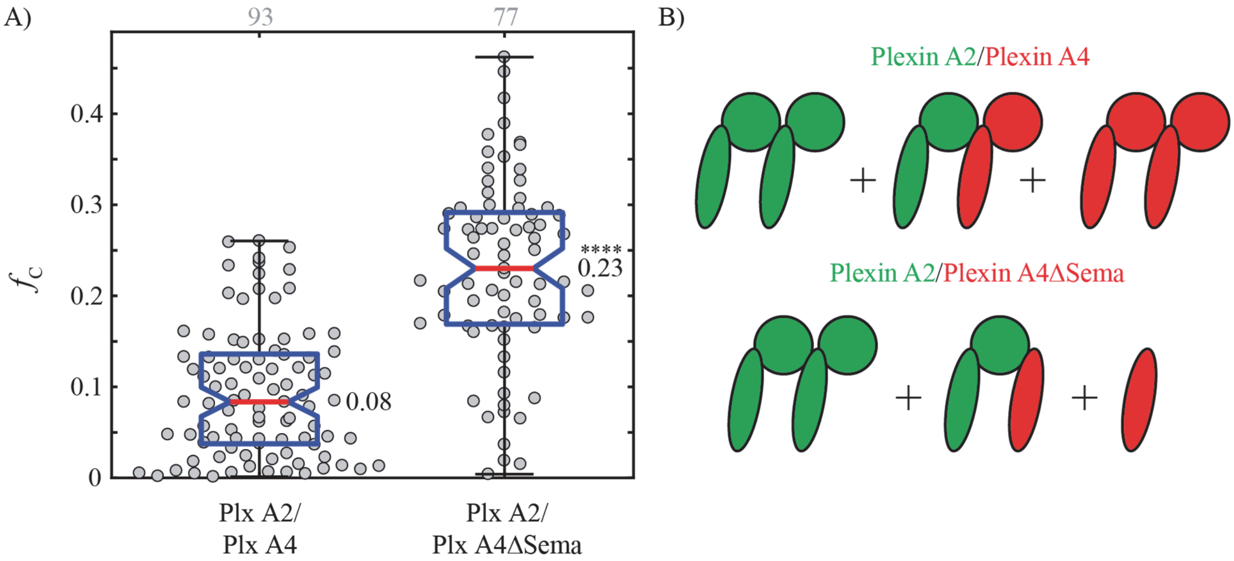


***Figure S3.*** *Fraction correlated of class A plexin mutant variants. A) Plexin A4 has the entire Sema domain deleted and shows a dramatic increase in correlation with WT Plexin A2. Each combination is significantly different from each other ($p<0.0001$). Grey numbers above each column represent the number of single cells analyzed. B) Diagram of receptor combinations to show how interactions would change due to mutation. Top, homotypic and heterotypic affinities likely allow equal mixing of dimers and cause an* $f_C$ *value associated with a weak dimer. Bottom, Removal of the Sema domain in Plexin A4 allows for a shift toward heterotypic interaction due to the lack of competition with homodimers of Plexin A4 leading to a drastic increase of the* $f_C$ *value.*

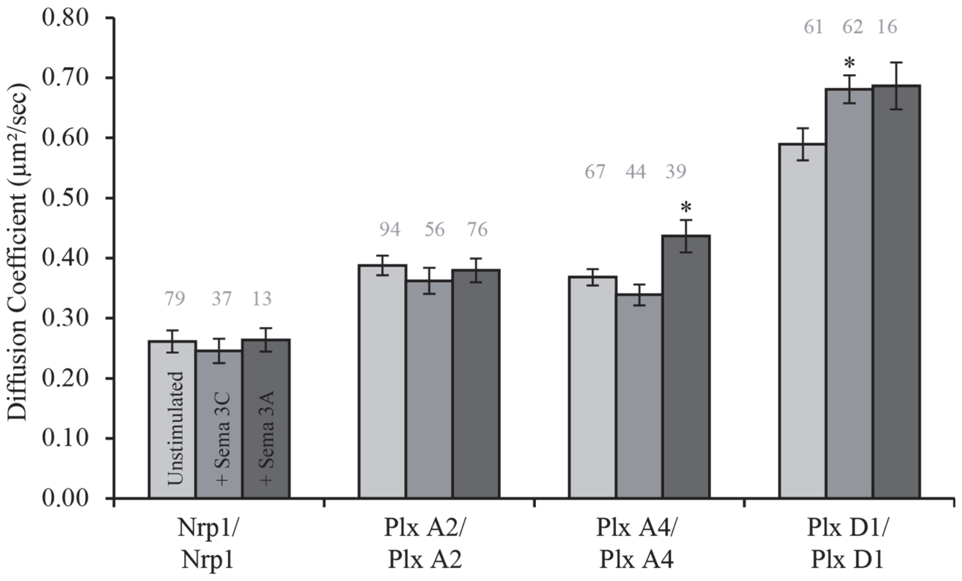


***Figure S4.*** *Diffusion coefficients for Nrp1 and Plexin A2 homodimers are unchanged following stimulation with either ligand, while both Plexin A4 and Plexin D1 are significantly increased ($p<0.05$) for one type of ligand stimulation.*

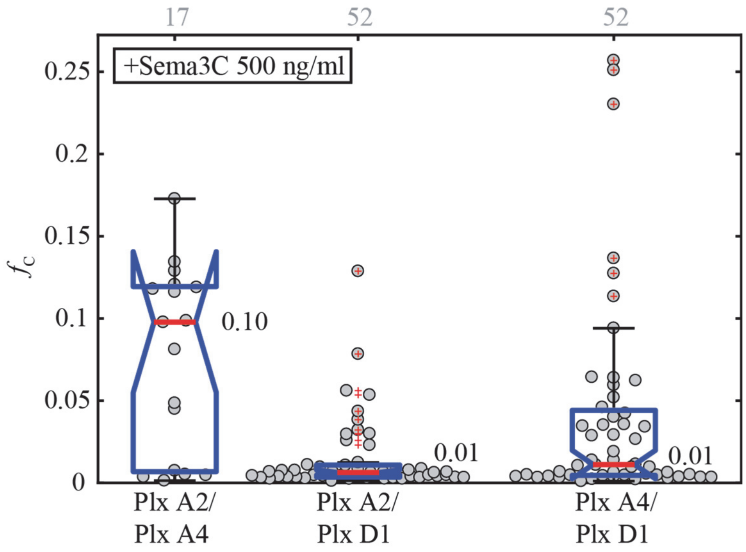


***Figure S5.*** *Heterotypic interaction of plexins following stimulation with Semaphorin 3C. Fraction correlated for each combination of plexin receptors shows each interaction is not significantly changed from the unstimulated group.*

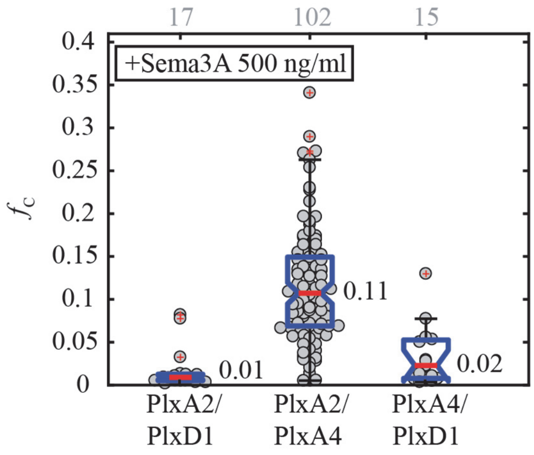


***Figure S6.*** *Heterotypic interaction of plexins following stimulation with Semaphorin 3A. Fraction correlated for each combination of plexin receptors shows each interaction is not significantly changed from the unstimulated group.*
